# Attractor dynamics reflect decision confidence in macaque prefrontal cortex

**DOI:** 10.1101/2023.09.17.558139

**Authors:** Siyu Wang, Rossella Falcone, Barry Richmond, Bruno B. Averbeck

**Affiliations:** Laboratory of Neuropsychology, National Institute of Mental Health, National Institutes of Health, Bethesda MD, 20892-4415; Leo M. Davidoff Department of Neurological Surgery, Albert Einstein College of Medicine Montefiore Medical Center, Bronx,NY, 10461

**Keywords:** population dynamics, decision certainty, prefrontal cortex, neurophysiology

## Abstract

Decisions are made with different degrees of consistency, and this consistency can be linked to the confidence that the best choice has been made. Theoretical work suggests that attractor dynamics in networks can account for choice consistency, but how this is implemented in the brain remains unclear. Here, we provide evidence that the energy landscape around attractor basins in population neural activity in prefrontal cortex reflects choice consistency. We trained two rhesus monkeys to make accept/reject decisions based on pretrained visual cues that signaled reward offers with different magnitudes and delays-to-reward. Monkeys made consistent decisions for very good and very bad offers, but decisions were less consistent for intermediate offers. Analysis of neural data showed that the attractor basins around patterns of activity reflecting decisions had steeper landscapes for offers that led to consistent decisions. Therefore, we provide neural evidence that energy landscapes predict decision consistency, which reflects decision confidence.

## INTRODUCTION

Decisions are made with varying levels of certainty. For example, we may be certain about which political candidate we prefer, but less certain about which entree to order at a restaurant. When we are certain about a choice, we are more likely to make that choice consistently. However, when we are uncertain, our choices are less consistent, and we may change our mind before making a final decision^1^. There is a substantial body of literature looking at theoretical, behavioral, and neural aspects of decision certainty, also referred to as decision confidence^2-4^. It has been shown that prefrontal and parietal cortical areas represent decision certainty^5-8^. While these studies have established that multiple brain areas contribute to decision certainty, they have not examined the network computational mechanisms that drive variability in decision making under different levels of certainty.

A prominent family of models that account for flexible decisions are the two-state attractor models. In these models, decisions are made when network activity settles into one of two attractor basins^9-15^. The models operationalize the idea that decision-related neural activity can be modeled as an object frictionally sliding on a landscape characterized by two valleys separated by a hill^16^. The valleys represent the attractor basins. The ball starts on top of the hill and is pushed into one of the valleys by evidence in favor of one of the choices. The decision corresponds to the valley in which the ball ends up^17^. These models account for a wide range of behavior including perceptual and value-based decision making^12,13^, and working memory^17^. Change-of-mind and variability in decision making is accounted for in these models by assuming that transient stimuli or noise pushes neural activity from one attractor basin (i.e., valley) to another^13,18-20^. Theoretical work using artificial network simulations suggests that choice consistency can be accounted for by the shape of the energy landscape around the attractor basins. When energy landscapes have steep sides and deep basins, corresponding to a high hill between the valleys, decisions will be made consistently, because it is hard to drive activity from one basin to another. However, when energy landscapes are relatively flat and attractor basins are shallow, decisions will be made less consistently, because it is easier to drive activity between attractor basins. It has been shown, for example, that reinforcement learning can increase the energy separation between the attractor basins, as optimal choices are learned, which can lead to increased consistency of choices^21,22^. When attractor basins are shallower, before values have been learned, choices are less consistent^21^.

Despite these advances in theory, less work has examined directly the dynamics in neural activity that underlies variability in decision making. Dynamical systems models have been used to understand neural activity in motor control^23^, decision making^24-26^, representation of head direction in rodent anterior thalamic nuclei^27^ and the encoding of aggression in the hypothalamus^28^. Most studies have focused on the average trajectories that underlie movements and choices as they unfold. Recently, however, researchers have examined flow fields and energy landscapes underlying these mean trajectories^27^ and explicitly examined attractor dynamics by fitting dynamical system models to neural data^26,28^. Studies on the computations underlying decision making have focused on evidence integration or drift-diffusion models^29,30^. These models have led to substantial insights, but they do not incorporate fixed-point dynamics, which may be an important component of neural decision mechanisms. To the best of our knowledge, there has been no direct neural evidence that the geometry of attractor basins, which reflect the energy landscape that drives neural activity, predicts decision certainty. In this work, we investigated population dynamics of prefrontal neurons using high channel-count recordings (768 channels; ^31^) while monkeys performed a decision-making task. We trained monkeys to choose between rejecting and accepting offers of different reward sizes and delay. By linking neural attractor dynamics to decision certainty, we provide evidence that the energy landscape in prefrontal cortex predicts decision certainty in behavior.

## RESULTS

We trained rhesus monkeys to perform a decision-making task^32^(Fig. 1A). In this task, animals had to reject or accept an offer by releasing a bar at different times. In each trial, animals were presented with one of nine possible reward offers, signaled by a visual stimulus. Each offer was a combination of reward size (2, 4 or 6 drops) and delay (1, 5 or 10 seconds; Fig. 1B). Monkeys began a trial by touching a bar and holding it for 500ms. Then a red square was presented at the center of the screen for 500ms.

**Figure 1.**
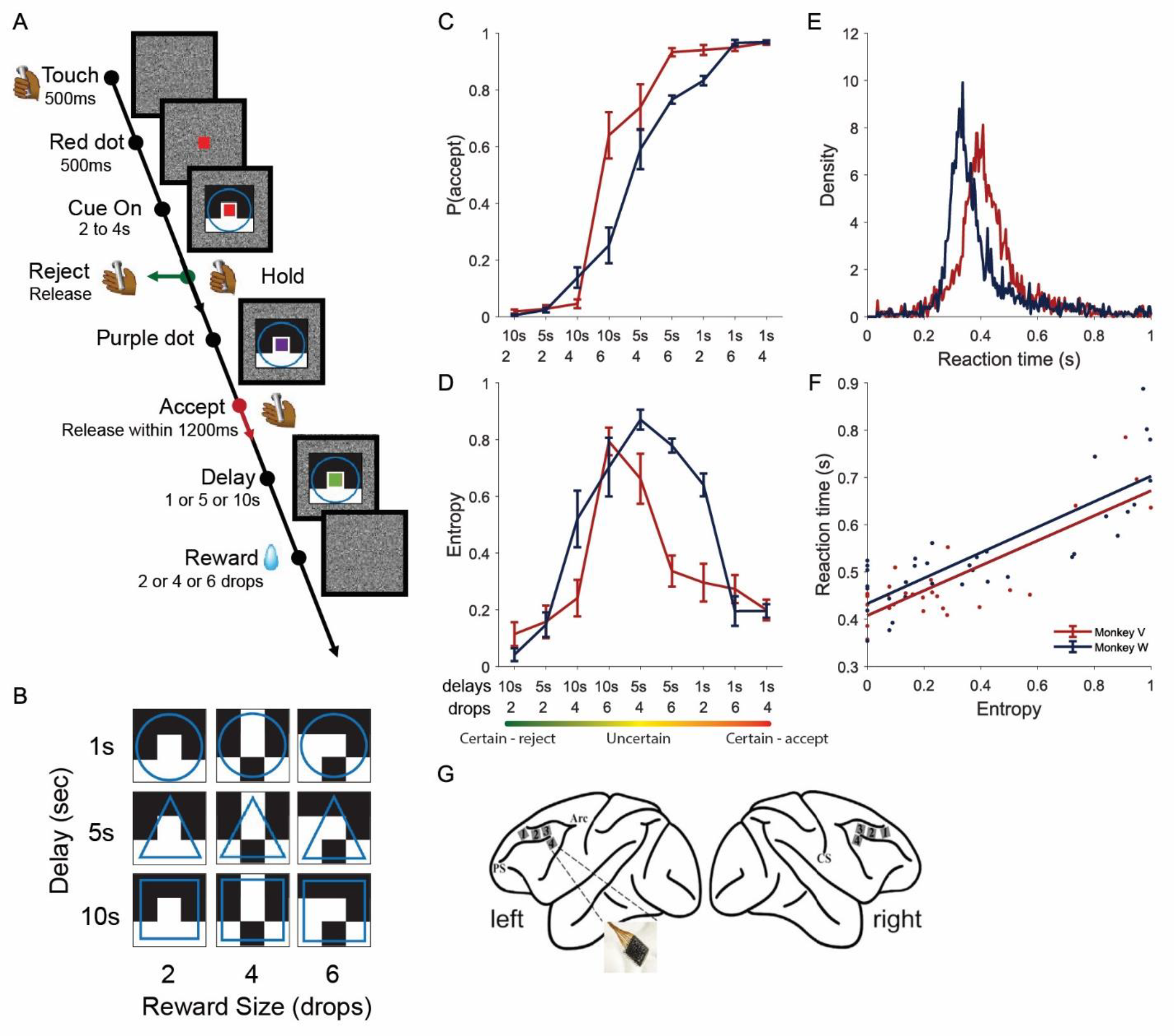
Behavioral task, behavior, and recording sites. A. Temporal-discounting task ^32^. In each trial, the animals first touched a bar, after which a red dot was presented. Next, a cue, which indicated the reward size (drops) and delay, was shown. If the animals released the bar before the red dot turned purple, the trial was considered rejected and, in the next trial, a new randomly selected cue was presented. If they held the bar until the red dot turned purple, and then released the bar between 200 and 1200 ms, the trial was considered accepted, and the indicated drops of liquid reward were delivered after the indicated delay. B. The nine visual cues used to indicate reward size and reward delay. C. Probability of accepting the given offer as a function of each offer of reward size and delay for the two monkeys. N = 8 for each animal. Data are presented as mean values +/-SEM. D. Entropy of p(accept) for each offer. N = 8 for each animal. Data are presented as mean values +/-SEM. Note entropy is high when offers are not consistently accepted or rejected, and low when offers are consistently accepted or rejected. E. Reaction time distributions for the two monkeys for reject trials only. F. Reaction time vs. decision entropy for reject trials. Note that individual points correspond to individual sessions. For monkey V, r = 0.82, p < 0.001; for monkey W, r = 0.82, p < 0.001. G. Recording locations in left and right hemispheres in each animal ^60^. PS: Principal Sulcus; Arc: Arcuate Sulcus; CS: Central Sulcus.

After that, one of the nine visual cues (Fig. 1B) was presented behind the central red square. The central square stayed red for a random period between 1500 and 3500ms before it turned purple. To reject the offer, the monkey had to release the bar before the square turned purple. After an offer was rejected, a new trial started, with a new cue randomly selected from the set of possible cues. To accept the offer, the monkey had to hold the bar until the red square turned purple and then release it between 200ms and 1200ms after the presentation of the purple square. After an offer was accepted, the purple square turned green, and the designated number of drops was delivered at the indicated delay. The visual cue was turned off during reward delivery. The monkeys completed on average 848 (Monkey V) and 761 (Monkey W) valid trials per session. Data from 16 sessions (8 per monkey) were analyzed.

We computed the proportion of times that monkeys accepted each offer, *p*(*accept*) (Fig. 1C). Monkeys almost always accepted offers with large rewards and short delays (e.g., 6 drops and 1s delay), and almost always rejected offers with small rewards and long delays (e.g., 2 drops and 10s delay). For other offers (e.g., 4 drops and a 5s delay), the acceptance probability was intermediate. The decision consistency for each offer was quantified by the entropy of choices (see Methods). Entropy is a natural measure of uncertainty in probability distributions^33^. Lower entropy characterized consistent choices for both the best and worst offers. Higher entropy characterized inconsistent choices for intermediate offers (Fig. 1D).

We also analyzed whether decision consistency was reflected in reaction time. Reaction times for reject decisions (from cue onset to bar release) peaked between 300 and 450ms for the two monkeys (Fig. 1E, Supplementary Fig. S1A). We found that the average reaction time correlated significantly with decision consistency for each offer in reject decisions (p < 0.001 for both monkeys, Fig. 1F). Reaction times were slower when decision consistency was smaller. This correlation may not entirely reflect decision consistency, but rather motivation. As a negative control, we analyzed response time for accept decisions (from purple dot to bar release). Accept response time also significantly correlated with decision consistency (p = 0.001 for monkey V, p < 0.001 for monkey W, Supplementary Fig. S1B). Since the decision, presumably, has already been made when the dot turns purple (Supplementary Fig. S1C), this correlation should not reflect the decision-making calculation. Rather it may reflect the speed of simple visual-motor reaction time. To control for the effect of motivation, we performed an additional combined regression analysis (see Methods) which included both p(accept), the probability of accepting an offer which serves as a measure of motivation, and choice entropy which serves as a measure of decision consistency. Our results showed that both p(accept) and choice entropy significantly predict reaction time for monkey W (p < 0.01 for p(accept), p < 0.05 for entropy), and only choice entropy statistically predict reaction time for monkey V (p > 0.05 for p(accept), p < 0.05 for entropy). This further suggests that decision consistency is reflected in reaction times, after controlling for motivation.

### Neurons in PFC show task-relevant activity

While the monkeys performed the behavioral task, we recorded the extracellular activity of neurons from prefrontal cortex using a bilateral implant of 8 multielectrode Utah-arrays (96 electrodes each), 4 in each hemisphere (Fig. 1G). We recorded on average 326.9 (Monkey V) and 527.5 (Monkey W) neurons simultaneously per session for the two monkeys. The recorded neurons were evenly distributed across left and right hemispheres.

We aligned the neural activity to the onset of the visual offer cues in each trial. Then we counted the spikes within a sliding window (50ms width, 50ms steps) from 1000ms before cue onset to 2000ms after cue onset for each trial and neuron. We performed a sliding window multivariate ANOVA analysis on spike counts (Fig. 2A, Supplementary Fig. S2). The results showed that a significant proportion of neurons responded to the offer features (reward size, delay, and their interaction) and the choice.

**Figure 2.**
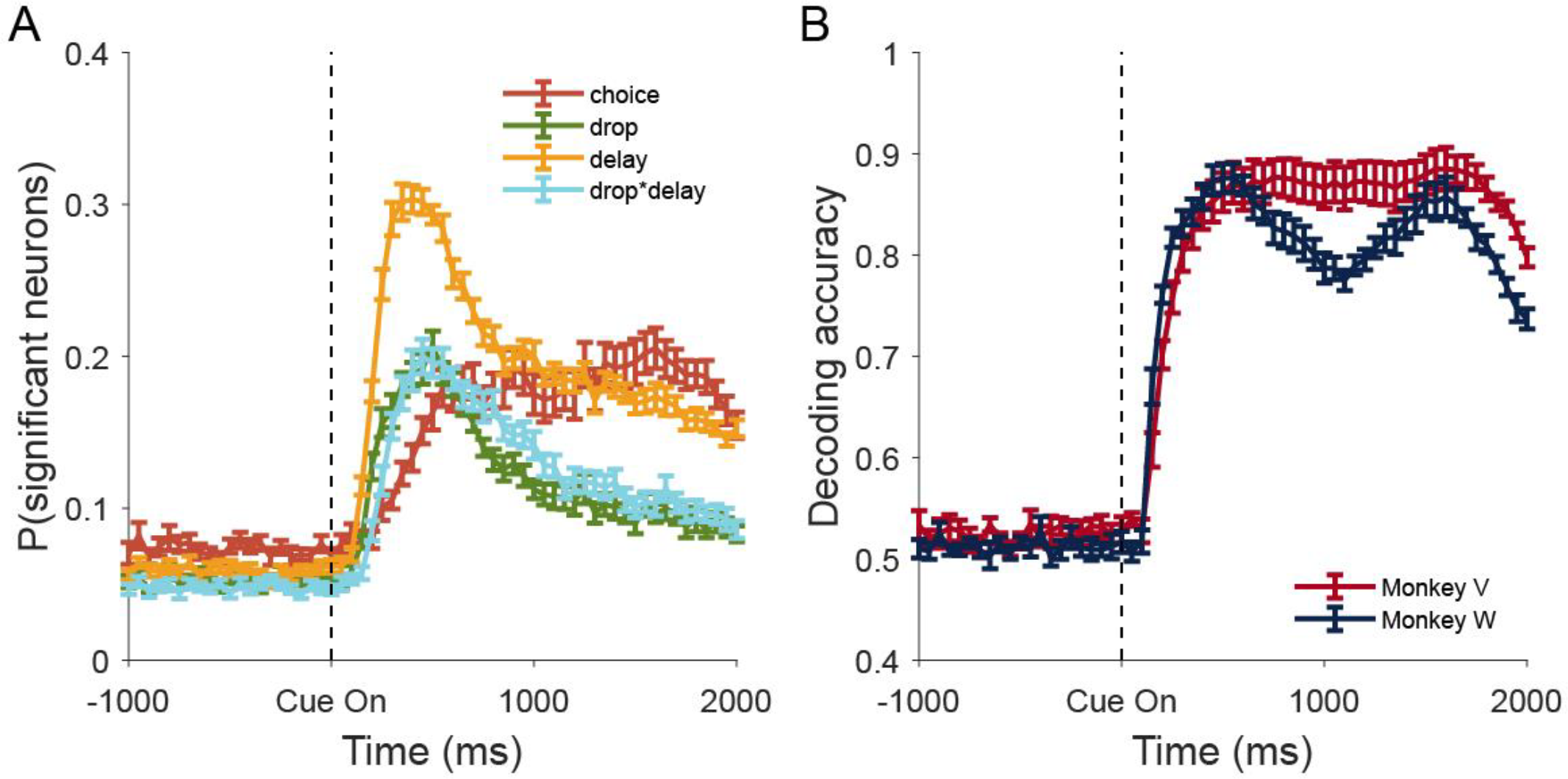
Neuron populations in PFC encode choice, and the offer value. Error bar indicates +/-SEM. N = 16 for both animals or n = 8 for each animal. A. Fraction of single neurons across both monkeys and recording sessions that significantly encoded each indicated task factor assessed with a multivariate ANOVA (F-test, alpha level set to 0.05). B. Decoding accuracy for accept vs. reject decisions from the 20-D PCA space defined for each session, as a function of time relative to cue onset.

Next, we carried out principal component analysis (PCA) on the simultaneously recorded neurons from each session. PCA was done on the mean spiking activity for each offer cue and each choice type (accept vs reject) in 50ms bins from 0 to 1000ms after cue onset. The first 20 principal components accounted for about 65% of the variance (Supplementary Fig. S3A). We could decode trial-by-trial choice (accept vs. reject), using a linear SVM with 10-fold cross validation, at close to 90% accuracy based on the neural population activity projected into the 20-dimensional PCA subspace (Fig. 2B). We were also able to decode information about reward size and delay from the 20-D neural subspace (Supplementary Fig. S3B, C). Adding more dimensions did not increase decoding performance (Supplementary Fig. S4). The decoding of the two monkeys’ choices were above baseline from 200ms after cue onset.

### Neural trajectories and energy landscapes in choice subspace

We examined the neural dynamics underlying decisions in a one-dimensional neural subspace using activity in the simultaneously recorded neurons from each session from each animal. This allowed us to visualize the way in which population activity, including trial-to-trial variability in population activity, tended to evolve as decisions developed. We began by reconstructing a numerical approximation to the energy landscape in this 1-D subspace (see Methods). To carry out this analysis we first projected the 20-dimensional population activity in each trial and each time bin, *P*_*t*_, onto the one-dimensional choice dimension defined by the SVM (Fig. 3A, B). This gave us a point in the 1-D space, *X*_*t*_, at each time t. We then computed time derivatives, *δ*_*t*_ = *X*_*t*+Δ*t*_ − *X*_*t*_, in the 1-D space (Fig. 3B). This was done for all times considered, and all trials. We binned the derivatives in the 1-D space, by finding all points *X*_*t*_ that fell into each spatial bin along the 1-D axis (Fig. 3C) and calculated the average derivative *δ*_*t*_ for all points starting at *X*_*t*_ that fell into each bin. This provided an estimate, on average, of how the neural activity evolved in this 1-D space, as a function of location in the 1-D space. These derivatives are estimates of the vector field for the dynamics underlying population activity (^34^, see Methods). The vector field characterizes the forces, driven by cortical networks and their inputs, acting on activity as it evolves through time.

**Figure 3.**
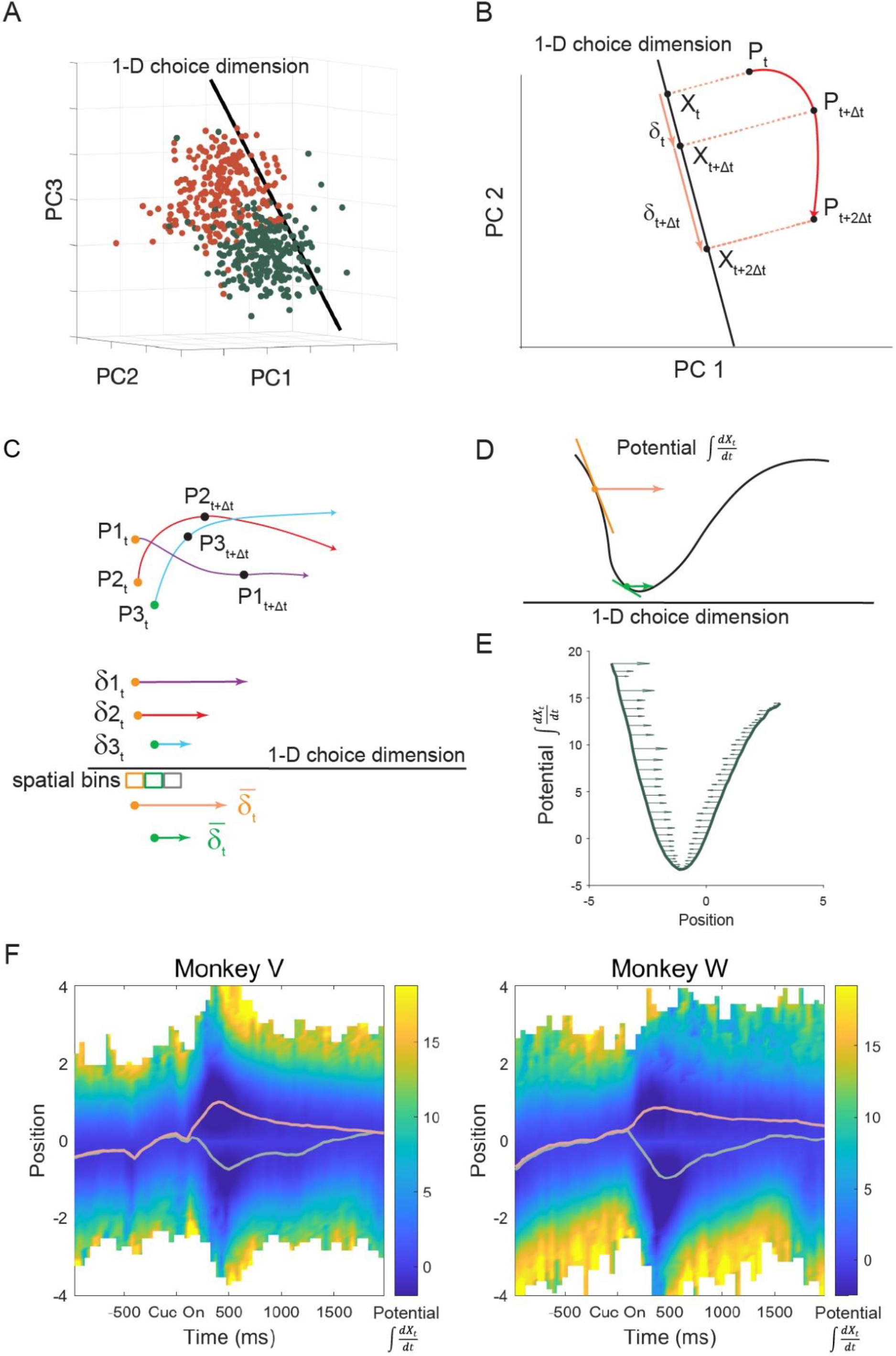
Reconstruction of energy landscape. A. The choice dimension defined by the SVM classifier, i.e., the dimension that is orthogonal to the separating hyperplane in the high-dimensional neural space (20-D PCA space). B. (cartoon) Activity, *P*_*t*_, in 50 ms time bins, in high-dimensional neural space is projected onto the 1-D choice dimension. This results in a time series of projected activity, *X*_*t*_. Activity was analyzed in 50 ms bins, so ∆*t* = 50 ms. Panel B illustrates the projected temporal trajectory of a single trial. C. (cartoon) Illustration of the temporal derivatives for 3 trials and the computation of mean derivatives in each spatial bin in the choice dimension. D. (cartoon) Integration over the mean derivatives in each spatial bin gives us the energy landscape or potential function. At each spatial location, the derivative of the energy landscape equals the mean derivative derived in C. E. The computed energy landscape for reject decisions at an example time point. F. The evolution of the energy landscapes, within the 1-D choice dimension, over time for both monkeys for accept and reject decisions. Position 0 is the SVM decision boundary and time 0 is cue onset. The light green line indicates the average activity for reject trials, and the red line indicates the average activity for accept trials. For the portion of the state space that was unvisited, there is no gradient or potential estimate, and it is left blank. The choice-dimension for all bins was the same and defined at the mean reaction time in each

In the next step we spatially integrated these average derivatives, across the 1-D space, to generate an estimate of the potential function that drove neural activity (Fig. 3D, illustration, Fig. 3E, data). On average, neural activity evolves from high potential energy to lower potential energy, similar to an object frictionally sliding on a landscape. Points with small spatial derivatives led to flat spots in the energy landscape, because when the neural activity fell into these locations it tended to stay close to these locations, on average. Correspondingly, points with large spatial derivatives led to steep spots on the energy landscape, because neural activity that fell into these locations tended to evolve away from them quickly, in the direction of decreasing energy (Fig. 3D). Stated another way, population activity tends to move from a high potential position to a low potential position, following the gradients.

We examined the energy landscapes computed separately for the accepted and rejected trials. For these analyses we averaged across cue conditions. We merged the landscapes at the SVM decision point (position 0), such that the landscape on one side of the decision point was derived from analyses of the accept trials, and the landscape on the other side was derived from analyses of the reject trials. By plotting these energy landscapes over time, our results showed that the attractor basins for accept and reject choices were deepest at around the time of the decision (∼300 to 600ms after cue onset) (Fig. 3F). We overlaid the average activity in this space, for accept and reject decisions, and it could be seen that the average followed the low point on the landscape across time (Fig. 3F).

While an energy landscape defined as the integral of the flow vectors can be computed in the 1-D subspace, true energy functions for higher dimensional neural subspaces may not exist. The existence of an energy function implies that the integration of gradients in the vector field between two points along different paths lead to the same quantity, or correspondingly that the integral along a closed path is zero. For 3 dimensions, the existence of an energy function requires the curl of the field (see Methods) to be zero at all points in space. Although we do not have access to the curl at all points in space, we empirically estimated the curl around the mean population trajectory based on our local estimate of the recurrent dynamics in a 3-dimensional neural subspace (see Methods). The estimated curl was not significantly different from zero at most time points (Supplementary Fig. S5). This does not prove the existence of an energy function. But we did not observe statistically significant non-zero curls. A non-zero curl would violate the assumption of the existence of an energy function.

### The geometry of attractor basins predicts choice consistency

We further investigated how the energy landscape in the 1-D subspace changed as a function of decision consistency. We tested the hypothesis that shallower basins at the time of decision were associated with lower decision consistency. During decision formation, neural activity will tend to frictionally slide downhill on these landscapes, towards the low points (attractor basins) that represent the decisions.

However, neural activity is also subject to noise, and the noise can drive the neural activity out of one basin and into another. Our hypothesis, therefore, suggests that deeper attractor basins will lead to more consistent decisions because variability in neural activity is less likely to stochastically drive population activity out of the deeper basins.

We computed the energy landscape separately for each of the nine cues, averaged across trials in which both accept and reject choices were made. These energy landscapes were computed for each session and averaged across sessions. We found that the landscapes had steeper basins following presentation of cues that had high decision consistency (accept decisions for large reward short delay, or rejection decisions for small reward long delay) and shallower attractor basins following cues that had low decision consistency (intermediate reward and delay) (Fig. 4).

**Figure 4.**
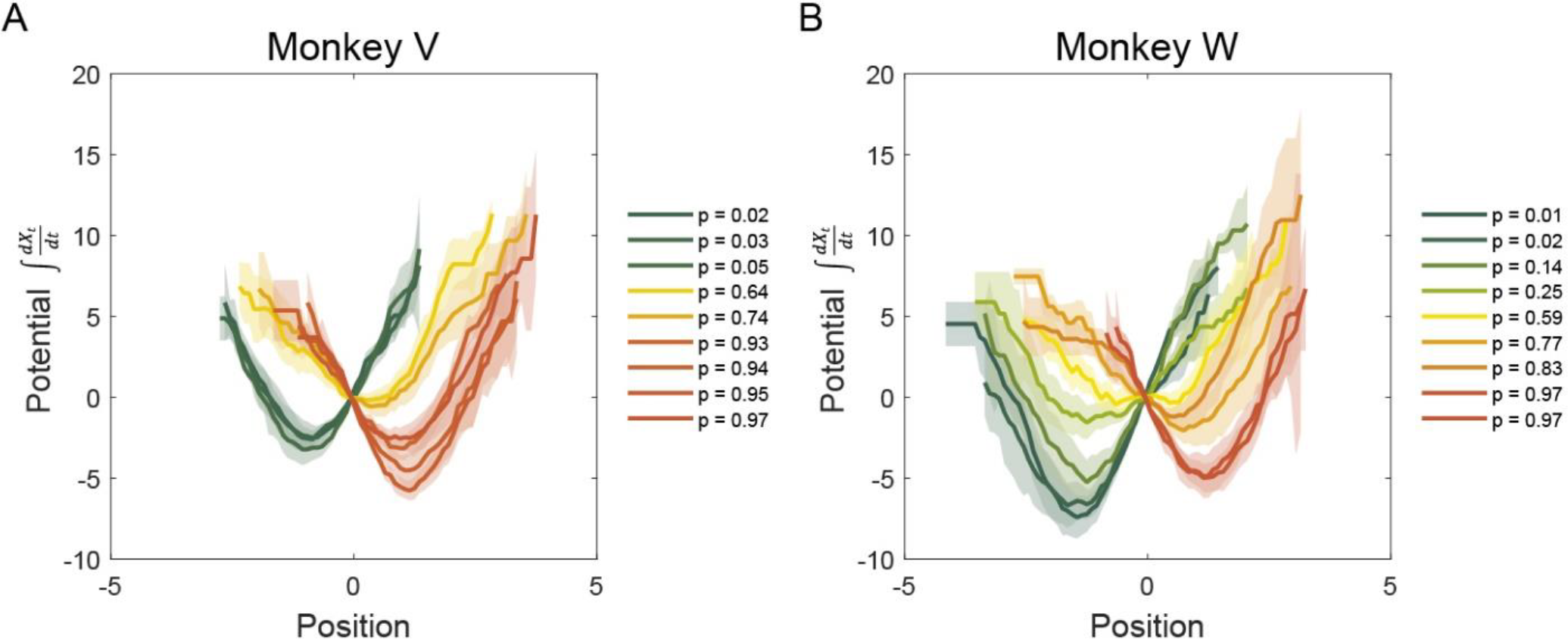
Numerical estimate of energy landscapes for each cue. Data are presented as mean values +/-SEM. A-B. The landscapes have qualitatively shallower attractor basins following presentation of cues that had low decision consistency for the two monkeys. The lines are separated by the probability of accepting the offer, and correspondingly, positive values in the choice dimensions reflect accept choices. We also examined landscapes separated by accept and reject decisions for the uncertain decisions (Extended Data Fig. 1).

Most of the landscapes appeared to have only one basin as opposed to two basins. This is partially because we have more data for the basin that corresponds to the more favorable choice by the animal for each cue. For example, the population activity rarely visits the negative portion of the 1-D subspace for cues which animals always choose to accept, thus we do not have an estimate of the reject basin. Furthermore, because our observation of population position is a noisy readout of the true position, this noise could potentially flatten the observed landscape. Partially because of this, there appears to be 1 basin even for cues that had low decision consistency in which both positive and negative parts of the subspace were visited. However, when we reconstructed the basins separately for these cues, for accept and reject choices, it could be seen that they had separate basins for each choice (Extended Data Fig. 1). We further examined this by comparing the fit of one and two basin models, and found that our data was better fitted by a two-basin model compared to a one-basin model (one-basin model: AIC = 2533.8; two-basin model: AIC = 2515.0). The minima of the reconstructed basins also shifted outward for consistent choices (Fig. 4), which was also reflected in the mean choice trajectories (Extended Data Fig. 2). These reconstructed landscapes are, therefore, consistent with our hypothesis. However, the differences across conditions could reflect differences in noise for the different conditions as well as the evidence in favor of each choice, as opposed to differences in attractor dynamics. Therefore, in the next analysis, we fit models to estimate the neural dynamics, as well as the effect of evidence, derived from a probabilistic model, and the overall noise level.

We fit dynamical systems models to compare three hypotheses that can account for the neural activity during decision formation. The simplest model assumes that activity in prefrontal cortex reflects only the decision, without reflecting either the evidence used to arrive at the decision, or neural dynamics. The second hypothesis is that the activity reflects a (strict) drift diffusion process. Drift-diffusion processes assume perfect integration of evidence, and no underlying fixed-point dynamics. Finally, we examined the hypothesis that choices are driven by both evidence accumulation and fixed-point dynamics. These three hypotheses can be compared by fitting a single model and examining the parameters. Specifically, we fit a model which predicted the time dependent neural activity as a function of a cue dependent evidence term, 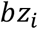 derived from the behavior (Extended Data Fig. 3A, also see Methods), and a cue and decision dependent dynamics term, *a*_*i*_(*X*_*t*_ − *x*_0,*j*_):

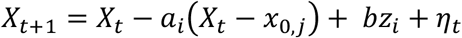

In this model *t* is time, *i* refers to one of the nine different cues and *j* indexes choice. The variables *a*_*i*_, *b*, and *x*_0,*j*_ are twelve free parameters, fit by nonlinear regression, to predict the time-dependent neural activity, *X*_*t*_, and *η*_*t*_ is zero-mean Gaussian white noise. The variable *a*_*i*_ characterizes the fixed-point dynamics. We refer to *a*_*i*_ as the retraction coefficient. A larger *a*_*i*_ reflects a steeper attractor basin. The variable *b* characterizes the strength of the evidence in favor of each choice, and the variable *x*_0,*j*_ characterizes the fixed-point of the undriven system for each choice (i.e., reject vs. accept). The undriven fixed point is an estimate of the mean position in state space to which the activity would go when the evidence term, *z*_*i*_, was 0 in the model. The evidence terms *z*_*i*_, were constants defined by fitting the choice consistency and reaction time behavioral data (Extended Data Fig. 3A). The variable *z*_*i*_ was estimated using three different versions of drift diffusion model-fitting methods. The results were similar between the three model-fits (Supplementary Fig. S6, S7). Note that if prefrontal activity represents a drift-diffusion process, then *a*_*i*_ = 0 and *b* > 0, and if the activity reflects only the choice without dynamics, then *a*_*i*_ = 1 and *b* = 0.

When examining the coefficients of the model, we found that the choice process, *x*_0,*j*_, was strongly represented around the reject decision reaction time and reflected the decision (Fig. 5A). Further, we found that the activity was affected by the evidence in favor of each choice, *z*_*i*_ (Fig. 5B). The evidence term was non-significant during the baseline hold, and peaked early, while the decision was being formed, but before the mean reaction time (Fig. 5B). Finally, we found that there were also significant fixed-point dynamics that did not reflect a perfect integration process, but that did reflect decision consistency (Fig. 5C). We further examined whether the depth of the attractor, characterized by the coefficients on the recurrent dynamics, *a*_*i*_, reflected choice entropy (Fig. 5D, E). We found that, at the time of choice, the dynamics were correlated with the choice entropy across cues (Fig. 5E). This correlation also was non-significant during the baseline hold, and became significant during choice formation, after the cues were shown (Fig. 5D). When the monkeys made consistent, low entropy decisions, either accept or reject, the retraction coefficients were larger (Fig. 5C). Thus, we found evidence for fixed-point dynamics in choice-related prefrontal cortex activity. We further found that the depth of the attractor basins reflected choice confidence, consistent with our numerical reconstruction of the energy landscape (Fig. 4) because a larger retraction coefficient reflects a steeper energy function. Thus, consistent decisions had deeper attractor basins, and inconsistent decisions had shallower basins.

**Figure 5.**
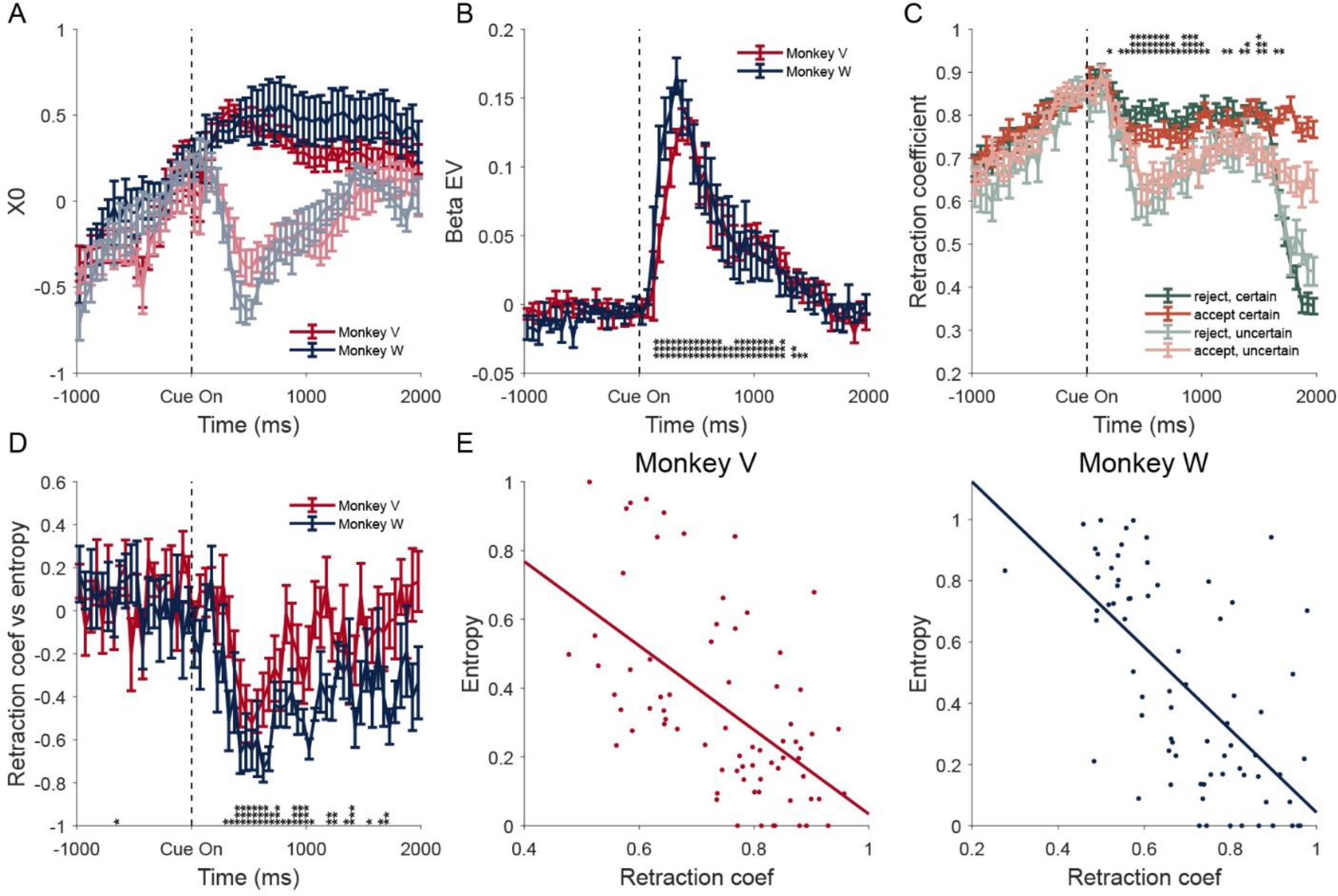
Neural activity reflects evidence accumulation and fixed-point dynamics. Error bars represent standard error of the mean. For plots that show data separately for each monkey, N = 8 as SEM was computed across sessions for each animal. A. Choice related activity, characterized as the undriven fixed points, *x*_0,*j*_, over time. Note the darker color curve is accept decision and lighter color curve is reject decision, for each monkey. B. Strength of evidence-driven neural activity over time, coefficient *b*. The curve peaks at 375 ms for monkey V and 325 ms for monkey W. A two-tailed one-sample t-test was performed at each time bin, uncorrected for multiple comparison. C. Retraction coefficients are higher for more certain decisions for both reject and accept choices. Certain accept = 90%+ vs. uncertain 50-90%. And certain reject 0-10% vs. uncertain 10-50% accept choices. Further, coefficients differ from both 0 and 1 and therefore there are fixed-point dynamics, and the system is not a perfect integrator. A one-tailed paired t-test was performed at each time bin between certain and uncertain conditions by first averaging between accept and reject conditions, uncorrected for multiple comparison. D. Correlation between retraction coefficient for each cue, and the behavioral entropy of that cue. A high retraction coefficient parameter corresponds to a steeper basin. A two-tailed one-sample t-test was performed at each time bin, uncorrected for multiple comparison. E. The retraction coefficients correlated significantly with decision consistency at the time of decision (i.e., mean reaction time in each session for each monkey). For monkey V, r = -0.56, p < 0.001; for monkey W, r = -0.64, p < 0.001. * = 0.05, ** = 0.01, *** = 0.001

We further examined the relationship between evidence accumulation and neural activity by fitting a variation of the model (Extended Data Fig. 4), in which we estimated a single constant, *h*_*i*_, for each cue, without explicitly including the evidence term, *z*_*i*_:

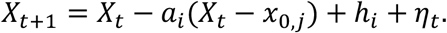

Thus, in this model we fit an evidence constant directly to the neural activity. We then compared the cue dependent constant, *h*_*i*_, fit to the neural data, to the cue dependent evidence term derived from the behavior, *z*_*i*_, and found that they correlated significantly, following onset of the cues (Extended Data Fig. 3B). This further supports the hypothesis that prefrontal neural activity reflects both evidence accumulation and fixed-point dynamics.

To exclude the possibility that the number of accept or reject choices for a particular cue drives the differences in estimates of retraction coefficients based on our model fitting procedure, we also conducted a control analysis. We refit the model after matching the number of trials between the consistent and inconsistent choice conditions. Our analysis suggests that the retraction coefficient was significantly larger even after matching the number of trials in the different conditions (Supplementary Fig. S8). We also controlled for the effect of ongoing change-of-mind on our model estimates. To consider changes-of-mind during the choice period, we predicted the current choice preference in each time bin (instead of using the final choice of the animal) and attributed the activity classified in this way to accept or reject decisions for the analysis. By repeating our linear dynamical system analysis using moment-by-moment estimates of choice, instead of the final choice, we accounted for the potential differences in the amount of “changes-of-mind” between different cue conditions. In this alternative analysis, our results still hold (Supplementary Fig. S9).

To further support our findings, we examined a 1-D choice dimension determined by an alternative classifier (see Methods, and Extended Data Fig. 5A-E). The estimated retraction coefficients, using choice dimensions defined by the SVM classifier vs the alternative classifier, were significantly correlated (Extended Data Fig. 5F). We also repeated our analysis in a 3-dimensional subspace spanned by the 1-D choice dimension and the first two principal components of the residual activity orthogonal to the choice dimension. Eigenvalues of the recurrent matrix were computed, in place of the 1-D retraction coefficients. Consistent with our 1-D analysis, we found similar correlations between choice entropy and the eigenvalues in the 3-D analysis. Larger eigenvalues, which reflect a steeper basin, were associated with low choice entropy and high choice consistency (Extended Data Fig. 6A-C). Moreover, the vector connecting the fitted undriven fixed points for accept and reject decisions in the 3-D analysis aligns with the 1-D choice dimension, and the cosine similarity between the two peaked at the time of choice (Extended Data Fig. 6D). Thus, analyses in 3-dimensions lead to results that agree with our findings in 1-dimension.

**Figure 6.**
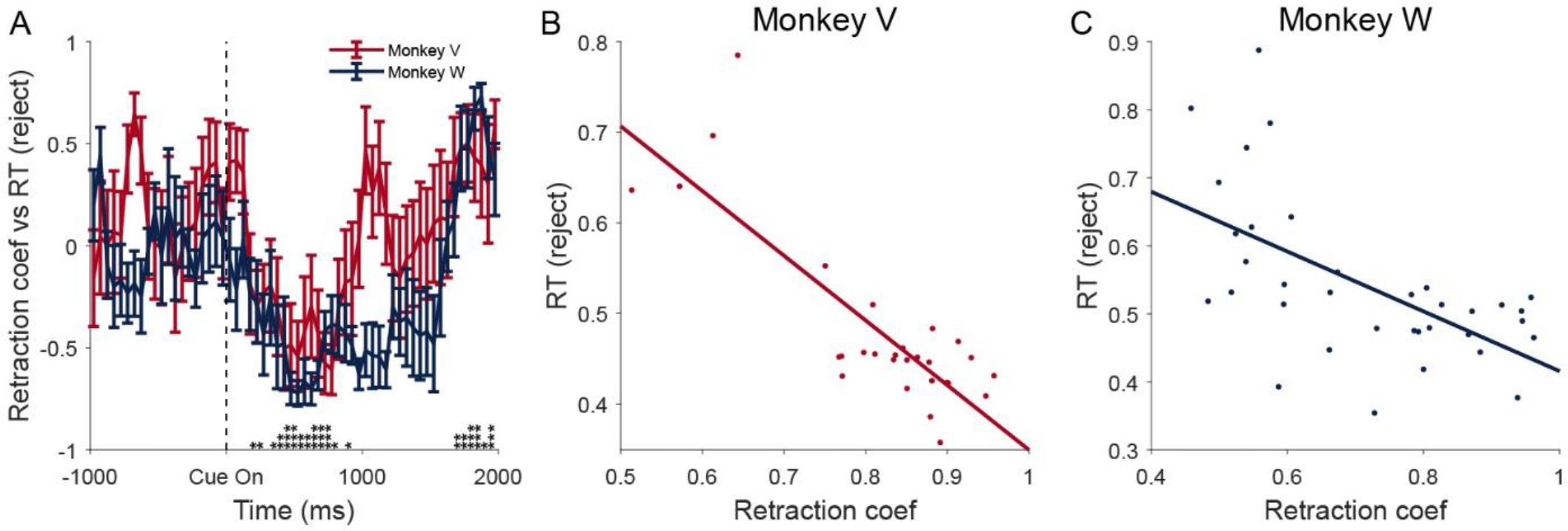
Retraction coefficients vs reaction time. For plots that show data separately for each monkey, N = 8 as SEM was computed across sessions for each animal. A. The correlations between retraction coefficients and reaction time for reject decisions over time. A two-tailed one-sample t-test was performed at each time bin, uncorrected for multiple comparison. B. The correlations between retraction coefficients at median time of reject decision time and reaction time for reject decisions (monkey V, r = -0.82, p < 0.001). C. Same as B for monkey W (monkey W, r = -0.59, p < 0.001).

We also examined how individual neurons’ responses to value affect the observed attractor dynamics. A separate sliding window multivariate ANOVA analysis with two features, choice and value, was carried out to examine individual neuron’s tuning to offer value. Behaviorally, offer value is reflected in the probability that the monkeys accept each offer. The results showed that the percentages of neurons that respond positively to value (accept probability) were 46.5% for Monkey V, and 37.8% for monkey W. Neurons that respond more to better offers also have higher loading weights on the 1-D choice dimension (Extended Data Fig. 7C). As a sanity check, we examined mean firing rates of individual neurons, and showed that negative preferring neurons have enhanced firing rates for reject offers, whereas positive preferring neurons have enhanced firing rates for accept offers (Extended Data Fig. 7A-B). Next, we repeated our dynamical system analysis using only positive, only negative or a mixture of positive and negative value neurons. For each session, the number of neurons for the three groups was selected to be the minimum between the number of positive and negative neurons to control for differences in population size. A random sample was taken for the other group and the mixture group. We observed that attractor retraction coefficients were larger for consistent offers for all three groups (Extended Data Fig. 8). This suggests that attractor dynamics do not differentially reflect activities of positive or negative offer preferring neurons.

### The steepness of attractor basins predicts reaction time

Since the derivatives in steeper basins are larger, we hypothesized that reaction times would also be smaller for steeper basins, under the assumption that choices are triggered when neural activity reaches a defined location in state space, which we refer to as the choice zone^11,23,35^. In theory, it takes fewer steps for neural activity to reach the choice zone in steeper basins with higher retraction coefficients.

Similar to the analysis of choice consistency vs retraction coefficients, we correlated the retraction coefficients with reaction time across cues. We found that the retraction coefficients significantly correlated with reaction time at the time of decision for reject choices in both monkeys (Fig. 6A-C).

As a negative control, we also computed the correlation between retraction coefficients extracted at the average reaction time for reject choices and response times for accept choices. Our results showed only a weak correlation between retraction coefficients and accept response time in Monkey W (p = 0.02), and none in Monkey V (p > 0.05) (Extended Data Fig. 9). This supports the suggestion that the correlation between reaction time for reject decisions and retraction coefficients is not merely due to the behavioral correlation between reaction time and decision consistency because the correlation is weaker for accept decisions. However, this only holds statistically in monkey V. Together, this supports our hypothesis that steeper attracter basins (higher *a*) are associated with shorter reaction times.

## DISCUSSION

In this paper, we investigated the neural underpinnings of choice consistency by analyzing dynamics in prefrontal population activity during decision making. We found that attractor basins were shallower following cues that signaled intermediate value offers, which also led to lower decision consistency.

Correspondingly, attractor basins were deeper following cues that signaled high or low value offers, that led to higher decision consistency. By fitting linear dynamical systems models to the neural data we found that the retraction coefficients, which characterize the energy landscape around the attractor, correlated significantly with both the entropy of choice and reaction time at the time of decision. Larger retraction coefficients, which are consistent with steeper energy landscapes, were associated with cues that elicited consistent decisions. Our results provide neural evidence that attractor dynamics predict decision consistency.

To visualize the dynamics of neural activity, we estimated the energy landscape in the state space of population activity. However, the existence of a true energy function is non-trivial and requires curls of the vector field to be zero (or in higher dimensions, the partial derivatives of the dynamics equation to be symmetric) at all spatial points in the neural state space. We estimated the curl around the mean population trajectory based on our fitted linear dynamical systems model in a 3-dimensional subspace. We did not observe curls statistically different from zero. However, this does not prove the existence of an energy function. First, failure to reject a null hypothesis does not prove the null hypothesis, thus failure to detect a non-zero curl does not prove the curl to be zero. Second, we only estimated the curl around the mean population trajectory and do not have global estimates of the curl at other spatial locations of the state space. Whether different neural systems, and different systems under different conditions, have energy functions is an interesting question. Considering whether networks in the brain have potential functions opens up several interesting neural and behavioral questions. Given plasticity rules, like spike-timing dependent plasticity, one wonders whether it is possible to ever return to the same brain state. Future studies are needed to directly examine the curl in different neural systems under different tasks.

We fit locally linear dynamical systems models, that let us compare three hypotheses, that can account for decision related activity in prefrontal cortex. The simplest hypothesis was that prefrontal activity would represent the choice, independent of the evidence that supported that choice, or dynamics related to the choice-formation process. The second hypothesis was that prefrontal activity would reflect a drift-diffusion process, that integrated evidence in favor of each choice. The third hypothesis was that prefrontal activity would reflect both evidence accumulation and fixed-point dynamics, consistent with network models that instantiate decision processes. We found that our dynamical systems model results supported the third hypothesis. Prefrontal activity reflected both the evidence in favor of each choice, as well as fixed-point dynamics. Furthermore, the dynamics were related to choice consistency, such that choices made with greater consistency were characterized by deeper attractor basins. The dynamical systems models also agreed with direct numerical reconstructions of the energy landscapes, which also reflected choice consistency. This association between energy landscapes and choice consistency has been predicted with computational models^19,21,22^. Intuitively it follows from the assumption that steeper landscapes will more likely lead to consistent decisions given the inherent stochasticity in neural activity. Variability in neural activity is less likely to drive neural population activity out of a deeper attractor basin. This association between choice consistency and the energy landscape has not been shown previously in neural activity, to our knowledge.

In our case, monkeys made accept and reject decisions based on the utility of an offer. Because reject decisions led to a random draw from the possible offers in the next trial, each trial’s offer could be compared to the average offer and the delay to the next trial. Consistent with previous work on this paradigm^32,36^, we found that offers were consistently accepted or rejected when the utilities were well above or below the average. Offers with intermediate values, however, were sometimes accepted and sometimes rejected. Because internal representations of offer values are stochastic, and choice probability is a function of value differences^6,37^, this leads to situations where choices are made with varying levels of consistency. Choice consistency can be related to decision certainty and decision confidence, which is the subjective probability that the chosen option has the highest value^3,5,8,38-40^.

Consistently accepted or rejected offers reflect high confidence choices, because the probability that the monkey is making the highest value decision is closer to 1, whereas inconsistently accepted or rejected offers reflect low confidence choices, because the probability that the monkey is making the highest value decision is closer to 0.5.

Several mechanisms have been proposed for representation of decision confidence. For example, drift-diffusion models (DDMs) have been developed to account for information integration in perceptual inference, value-based decision making, and memory recall tasks, and used to model decision confidence^6,38,41^. In perceptual inference, it is assumed that an internal process is integrating noisy information from an external stimulus^38^. In value-based decision making it is assumed that an internal process is integrating a noisy internal representation of offer value^6,37^. Across these models, the integration process can be related to a Bayesian posterior probability that a given choice is correct, and these posteriors can be used to define decision confidence. These models, and the associated Bayesian posteriors, provide good descriptions of the formal mechanism that gives rise to confidence, in many^42^, but not all^43,44^ conditions. However, they do not specify the neural mechanisms by which the brain implements these formal computations.

Network models for perceptual and value-based decision making have also been developed^10-12,14,18,20,45^. These models reproduce changes in mean population activity with decision confidence^14,45^. They do this by integrating activity from pools of neurons representing chosen and unchosen options. Because neural activity levels are higher for options when they are chosen with high confidence, confidence can be read out by integrating across the chosen and unchosen option pools. This suggests a mechanism by which activity levels related to certainty about chosen options^14^ can be converted directly to choice-independent confidence estimates^8^. Changes in the energy landscape of most of these networks, when choices are driven by different levels of information, have not been explored. Recent work, however, has examined changes in neural dynamics when reinforcement learning drives decisions from uncertain to more certain^21^. This study found that before learning which of two option was more valuable, in an RL task^46^, choice options were not well separated across the energy landscape. However, after learning, the options were better separated.

In empirical work, previous studies have shown that certainty, or probability distributions over sensory variables, can be estimated by decoding probabilistic population codes^42,47,48^, or through related sampling models^49^. When choice is decoded from neural activity, posterior probabilities of choices have also been shown to reflect decision consistency, and therefore average decision confidence^3,5,50^. Single neurons in OFC have been shown to directly encode decision confidence, independent of chosen actions, in odor and auditory discrimination tasks^8,51^. Thus, mean activity levels have been shown to reflect decision confidence.

We also found that the fixed-point of the dynamics, related to the mean choice-related activity, associated with options that were chosen more consistently was represented farther from the decision boundary. The fixed-point of the dynamics is given by 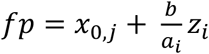 Therefore, if we know the fixed point of the undriven (*z*_*i*_ = 0) dynamics, *x*_0,*j*_, decision confidence can be read out by observing the fixed-point into which the system settles, at the decision time. (Note that *a*_*i*_ also scales with *z*_*i*_ and therefore *a*_*i*_ has to scale sub-linearly for this to hold.) This distinguishes the dynamical representation of evidence, from a DDM with a bound, because activity level (the position of the diffusion particle) and time have to be available to estimate the Bayesian posterior in a DDM. Furthermore, in our model activity settles into a fixed point that represents the evidence in a time-independent way. In a DDM, activity continues to increase over time, unless one assumes a sticky bound, and therefore it never directly represents the evidence. Whether perceptual decision-making processes would also show dynamics similar to what we have seen in the context of value-based decision making is an open question.

Because the fixed point reflects the evidence, confidence would not have to be estimated from the energy landscape, i.e., one would not have to compute an estimate of the energy landscape on a single trial to estimate confidence, even though the dynamics are correlated with decision confidence. The dynamics, however, reflect the energy required to change a decision. Flatter dynamics reflect the fact that choices are less consistent when they are associated with cues that predict intermediate value offers. The dynamics we have observed, therefore, likely reflect the changeability of a deliberative choice process when cues represent intermediate value offers, or a commitment to a decision when cues represent very good or very bad offers. Due to the correlational nature of our analysis, future perturbation studies are required to examine the causal role of attractor dynamics in change-of-mind and decision confidence. Related dynamics may also underlie observed variability in population neural choice representations seen in other decision-making tasks^52,53^. Because dynamics are also correlated with mean distances in decision space, this also raises the question of whether dynamics and posterior probabilities are linked, in an obligatory way, in biological and artificial networks.

## Conclusion

There is a substantial body of theoretical work that studies how computations in networks can underlie complex behavior^12,16,27,54-58^. However, there has been less experimental progress in this area, except in motor control^59^ and hippocampal circuitry^27,58^. In the present study, we have begun to make progress on linking computations in networks to population representations during decision making. We found that neural dynamics reflected choice consistency and reaction times, when monkeys accepted and rejected offers with different values. The neural dynamics for consistent decisions were characterized by deep attractor basins and large retraction coefficients in a dynamical systems model. The neural dynamics for inconsistent decisions had shallower attractor basins and correspondingly smaller retraction coefficients. Thus, the dynamic landscape of population neural activity in prefrontal cortex reflects choice commitment during value-based decision making. These results extend the insights into behavior that can be obtained by studying populations of simultaneously recorded neurons, and directly link theoretical work on network computations to experimental results.

## Acknowledgements

This work was supported by the intramural research program of NIMH (ZIA MH002928 BA). The authors thank Cary Robinson and Yi Wei for their assistance in spike sorting.

## Author Contributions

R.F. and B.R. designed the behavioral task. R.F. collected the data. S.W. and B.A. developed the analytical approach. S.W. analyzed the data. S.W. and B.A. wrote the manuscript, with input from R.F. and B.R.

## METHODS

### Animals

All experimental procedures were performed in accordance with the ILAR Guide for the Care and Use of Laboratory Animals and were approved by the Animal Care and Use Committee of the National Institute of Mental Health. Two male monkeys (Macaca mulatta, W – 6.5 kg, age 4.5yo, V – 7.5 kg, age 5yo) were used as subjects in this study. No statistical methods were used to pre-determine sample sizes, but our sample size of two monkeys is the field standard.

### Decision-making task

Monkeys were trained to perform a decision-making task while they were seated in front of a computer screen. In this task, animals had to choose to either reject or accept a reward offer that differs in reward size (drops) and delay time. Different reward offers were indicated by different pretrained visual cues that indicate different combinations of reward size (2, 4 or 6 drops) and delay time (1, 5 or 10s). In each trial, one of the 9 visual cues was pseudo-randomly selected and presented.

On each trial, animals had to first touch the bar and hold it for 500ms. Next, a small red square was presented at the center of the screen for 500ms. After that, one of the 9 visual cues was presented behind the red square at the center of the screen. The center square stayed red for a discrete random period at one of the five levels (1.5, 2, 2.5, 3 or 3.5 seconds) before it became purple. Animals reported their decisions by either releasing the bar anytime during the red period to reject the current offer or between 200 and 1200 ms after the purple square was presented to accept the offer. If the animal rejected the offer, a new trial started immediately. If the animal accepted the offer, the purple square became green and liquid reward of the indicated size was delivered after the indicated delay. The visual cue was turned off during reward delivery. If the monkeys released the bar before cue onset, within 200ms of presentation of the purple square onset or if they never released the bar during the trial, an error cue appeared on the screen for 1000 ms and the trial was considered an error. After an error trial the same cue was presented in the next trial. Error trials from both animals were excluded from the analysis.

### Data collection

A noncommercial software package, REX (version 8.0, ^61^), was used to control stimuli in the decision-making task. A commercial software package, Ripple Grapevine (version 3.2, https://rippleneuro.com/ripple-products/grapevine-processors/), was used for neural recordings.

Custom software (WangSpikeSorter; version 1.0) was used to perform offline spike sorting. Neurons that did not significantly predict any task related feature (reward size, delay, choice) were excluded from the analysis. Data collection was performed blind to the conditions of the experiments.

### Computation of choice consistency and decision evidence

Choice consistency is quantified by the entropy of choices. The entropy of choices is defined as the entropy of a Bernoulli process in which binary choices are generated with probability *p* and 1 − *p*. Specifically, the choice entropy can be written as a function *H*(*p*) of the probability that animals accept the offer, *p*.

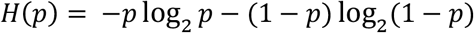

Decision evidence on the other hand is estimated by the drift rate in the drift diffusion model. The amount of evidence for choice of option *i* at time *t* can be estimated by ^62^:

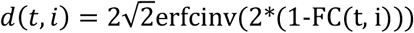

FC is the fraction of correct (preferred for that cue) choices, in this case *FC* = max (*p*, 1 − *p*).

The drift rate for each cue, *i*, was then estimated as

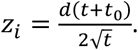

Where, *t*_0_ is a non-decision time and *t* is the reaction time minus the non-decision time. The non-decision time is not immediately available in this task. However, previous work has shown that non-decision time is under 200 ms. Therefore, we calculated evidence under a series of non-decision times (0, 50, 100, 150, and 200ms), and examined correlations in evidence, across cues, for different values of non-decision time. For the extreme values (0 vs 200ms), the calculated drift rates are correlated at R = 0.99, p < 0.01. Since estimated drift rates using different non-decision-times are strongly correlated, we adopted the average of the 5 sets of estimated drift rates as the final estimate for drift rates.

The accept decision times are not immediately available, since animals make accept decisions by not releasing the bar when the cue is presented, until the center red square turns purple which happens 1.5-3.5s after cue onset. Note that accept decision time is different from the accept response time analyzed in Supplementary Fig. 1, which instead measures the bar release reaction time after the red dot turns purple. Therefore, we inferred the accept decision times using the regression between entropy and reaction time for reject choices, and then using the entropy for accept choices. We carried out this approximation in two ways. First, we fit a linear model between reject RT and entropy for each session and estimated the accept decision time based on the session-specific regression model. Second, we fit a linear model between reject RT and entropy by pooling data from all sessions and estimate the accept decision time based on the pooled regression model. Results using either approach were similar.

In addition to using closed-form formulas to calculate drift rates as stated above, we also directly fit two variants of the drift diffusion model (DDM), vanilla DDM and DDM with collapsing bound, directly to the trial-by-trial reaction time and choice data. Since we only have access to trial-by-trial reaction time for reject trials, analysis was restricted to reject trials. Model-fitting was carried out using a recently published python library pyDDM (version 0.6.1) ^63^. Drift rates estimated using all three methods (closed-form solution, vanilla DDM, and DDM with collapsing bound) significantly correlate with each other in both animals (Supplementary Fig. S6).

For correlation of reaction time and choice-entropy, the reaction times were z-scored within each session. For plotting purposes, the z-scored reaction times were transformed to the overall reaction times, by multiplying by the overall average (i.e., across session) standard deviation and adding the overall average reaction time. This transformation does not affect the statistics, because all values were multiple by the same constant, and the same constant was added to all values.

For the combined reaction time regression, we have

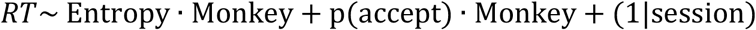

Both interactions between entropy and monkey, and between p(accept) and monkey were included in the model. A random effect of session was included to account for variations of RTs across sessions. The dependent variable here is reaction time, for both accept trials and reject trials. Reaction time for reject trial is the interval between cue onset and bar release while the square is red. Reaction time for accept trials, on the other hand, measures the response time between when the red square turns purple and bar release.

### Analysis of single unit responses

ANOVAs were applied to the single unit data to assess response association in sliding windows (50ms bin, 50ms steps) time locked to offer cue onset. The dependent variable was the spike count in each time bin. The ANOVA included main effects of choice, delay time, reward size, and interactions between delay time and reward size. A separate ANOVA was carried out which included the main effects of cue accept probability and choice. Data distribution was assumed to be normal but this was not formally tested.

### Dimensionality reduction of population activity and decoding analysis

All population analyses were conducted on simultaneously recorded neurons from a single session from a single animal. We performed dimensionality reduction using Principal Component Analysis. The mean spiking activity of each neuron was computed for each visual cue and each choice in 50 ms bins from 0 to 1000 ms after cue onset. This led to 18 time series of mean firing activity for each neuron. The time-series for the different conditions were then appended into one long vector for each cell, and the covariance matrix across simultaneously recorded neurons from a single session was then computed.

The first 20 principal components of the covariance matrix were kept for further analysis as they explained above 60% of the total variance. Including additional dimensions did not increase decoding performance for predicting monkeys’ choices (Supplementary Fig. S4).

A linear Support Vector Machine classifier was used to predict trial-by-trial variations in three task variables (choice, delay time and reward size) separately based on the first 20 principal component scores at each time bin. We used a 10-fold cross validation to evaluate the accuracy of the classifier.

In order to estimate and visualize the attractor dynamics of the population activity, we further projected the 20-dimensional neural data (based on Principal Component Analysis as described above) onto a single dimension, the choice dimension. The choice dimension was defined by the Support Vector Machine classifier and was the dimension perpendicular to the separating hyperplane in the SVM classifier. An alternative way of defining the choice was by computing the difference between the average population activity vector in the 20-dimensional subspace for accept vs reject trials. Results using this alternative definition were included in Extended Data Fig. 5.

### Estimation of the energy landscape in the 1-dimensional neural subspace

We reconstructed the energy landscape by numerically estimating the flow field in a 1-D space, and then spatially integrating the flow field^34^. Thus, we assume the neural dynamics of the population activity *X*_*t*_, are governed by a first-order (at this point potentially nonlinear) system,

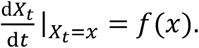

And then defined the spatial derivative of the potential function *V*(*x*),

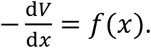

For a first order system 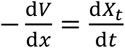. Therefore, the potential function, *V*(*x*), which we assume exists, is given by the spatial integral of the time derivative.

To estimate the time derivative, we defined the population activity in the 1-D subspace at time *t* and trial *i* as 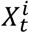 We first computed the expected value, ***E***, of the time derivative of population activity *X*_*t*_ at binned spatial locations [*x, x* + Δ*x*] and time *t*, with the expectation taken over trials, *i*.

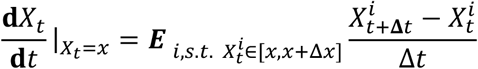

The interval Δ*t* = 50*ms*. Then we took the spatial integral over these time derivatives to get the potential function *V*(*x, t*) at location *x* and time *t*,

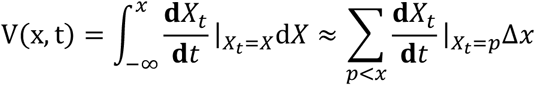

The integral results in an arbitrary constant. Potentials were always set to 0 at the center of the 1-D decision space.

### Linear dynamics analysis

Furthermore, we fit a linear dynamical system model to estimate the retraction coefficient ***a*** for each reward offer. Here we assume that in the low-D space(s) the dynamics could be approximated as a first-order linear system in which we model both fixed-point dynamics and evidence input to the system.

This gives the following equation:

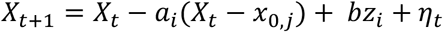

Here *X*_*t*_ is the activity at time *t*, and *η*_*t*_ is zero-mean Gaussian white noise. The coefficients *a*_*i*_ characterize the depth of the attractor, with larger values corresponding to deeper attractors. The coefficient *b* characterizes the strength of the evidence, and *x*_0,*j*_ is the fixed point of the undriven dynamics (*z*_*i*_ = 0) which also corresponds to the choice-related component of the fixed-point.

This model was fit to activity projected into the 1-D choice dimension, averaged in 50 ms bins. Thus, the activity in a 50 ms bin, *X*_*t*_ was used to predict the activity in the next 50 ms bin, *X*_*t*+1_. The model was fit to data from all correctly executed trials. From this regression, the retraction coefficient *a*_*i*_ was extracted.

Parameter recovery analysis of the linear dynamical model suggests that our model can be robustly fit to the data (Supplementary Fig. S10). We also performed posterior predictive checks to see if our model could qualitatively capture the differences in energy landscape that we observed in Fig. 4. In this analysis, we start with the recorded neural activity at the -1000ms bin, then simulated bin-by-bin the population activity in the next time bin using our fitted model. The noise term in the model is sampled from the empirical distribution of residuals from the model. Our model could indeed qualitatively capture the data (Extended Data Fig. 10).

We also fit a variant of the equation given as:

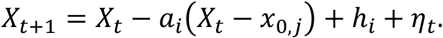

In this equation, *h*_*i*_ is a cue-specific term reflecting evidence accumulation. We fit *h*_*i*_ as a constant, and then subsequently correlated *h*_*i*_ with *z*_*i*_.

### Multi-dimensional linear dynamics analysis and calculation of curls

A multi-dimensional version of the linear dynamics analysis was also estimated in a 3-D subspace. A principal component analysis was performed on the residual neural activity orthogonal to our 1-D choice dimension. The 3-D subspace was then composed of the choice dimension and the first two principal components of the residual activity.

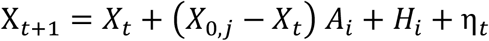

Here *X*_*t*+1_ and *X*_*t*_ are 3 dimensional vectors and represent the population activity at time *t*. The 3 x 3 matrix *A*_*i*_ is the recurrent matrix. *X*_0,*j*_ is the fixed point for choice *j. H*_*i*_ is the constant drift for cue *i. η*_*t*_ is zero-mean Gaussian white noise. Because the dynamics were non-stationary, the model was fit to a moving window of 3 time bins from each trial. The window was advanced by 50 ms, and 3 bins of 50 ms were fit, etc. Thus, for each time window, we used 3 consecutive time bins from each trial, and all valid trials, for one regression fit. This gave us temporal resolution on the change in dynamics, while also providing pooling over time within each trial (3 time bins) to increase statistical power.

Eigen values of the recurrent matrices *A*_*i*_ were computed for each cue *i*. Correlations between the eigenvalues and choice entropy were computed.

The 3-dimensional curl around the population activity *X*_*t*_ in the 3-D subspace was computed as (*A*_23_ − *A*_32_, *A*_13_ − *A*_31_, *A*_12_ − *A*_21_). Here *A*_*ij*_ represents the element of the recurrent matrix *A* at the *i*^*th*^ row and *j*^*th*^ column.

## Data Availability

Data from the manuscript can be found in the following figshare repository.

Wang, Siyu; Falcone, Rossella; J. Richmond, Barry; Averbeck, Bruno (2023). data for “Attractor dynamics reflect decision confidence in macaque prefrontal cortex”. figshare. Dataset. https://doi.org/10.6084/m9.figshare.21701282

## Code Availability

Custom spike sorting and data analysis codes were used. Codes for the custom spike sorter can be found at https://github.com/wangxsiyu/WangGit_Pilot_SpikeSorter.git. Custom MATLAB scripts used to perform all analysis and generate all figures can be found at https://github.com/wangxsiyu/Paper_NN-A81142A.git

